# Functional interplay between arginyl-tRNA synthetases and arginyltransferase

**DOI:** 10.1101/2022.01.11.475907

**Authors:** Irem Avcilar-Kucukgoze, Anna Kashina

## Abstract

Protein arginylation, mediated by arginyltransferase ATE1, is a posttranslational modification of emerging biological importance that consists of transfer of the amino acid Arg to protein and peptide substrates. ATE1 utilizes charged tRNA^Arg^ as the donor of the arginyl group, which depends on the activity of Arg-tRNA synthetases (RARS) and is also utilized in translation. The mechanisms that regulate the functional balance between ATE1, RARS and translation are unknown. Here we addressed the functional interplay between these mechanisms using intracellular arginylation sensor in cell lines with overexpression or deletion of ATE1 and RARS isoforms. We find that arginylation levels depend on the physiological state of the cells but are not directly affected by translation activity or availability of RARS isoforms. However, displacement of RARS from the multi-synthetase complex leads to an increase in intracellular arginylation independently of RARS enzymatic activity. This effect is accompanied by ATE1’s redistribution into the cytosol. Our results provide the first comprehensive analysis of the interdependence between translation, arginyl-tRNA synthesis, and arginylation.

**Key Points:**

1. Intracellular arginylation depends on the physiological state of the cell, but does not compete with the translation machinery
2. A fraction of ATE1 binds directly to both long and short Arg-tRNA synthetases (RARS)
3. Displacement of long RARS from the multi-tRNA synthetase complex increases cytosolic fraction and activity of ATE1

## Introduction

Protein arginylation, mediated by arginyltransferase ATE1, is a biological regulatory mechanism that involves addition of the amino acid Arg to proteins and peptides [1]. Over a hundred of in vivo arginylation targets have been identified [2-7], implicating ATE1 in regulation of a diversity of key cellular and organismal processes including cell migration [8], nucleotide biosynthesis [9], neurodegeneration [2], and cancer [10]. ATE1 is encoded by a single gene in animals and fungi and by two genes in plants. In higher vertebrates *Ate1* gene generates four alternatively spliced isoforms [11]. It is unclear how these four highly similar enzymes can perform a multitude of diverse in vivo functions, and what intracellular factors contribute to this regulation. To date, very few ATE1 functional partners have been identified, and very little is known about the intracellular mechanisms that balance arginylation with other pathways that utilize the same molecules as substrates, such as protein synthesis.

ATE1 requires Arg conjugated to tRNA^Arg^ as the donor of the arginyl group, and thus, arginylation directly depends on the activity of arginyl-tRNA synthetases (RARS). In principle, this dependence puts arginylation into direct competition with translation, which also depends on RARS, along with other aminoacyl-tRNA synthetases (AARSs), to provide aminoacyl-tRNA (aa-tRNA) for the growing polypeptide chains. Previous work from our lab showed that, in addition to full-length tRNA^Arg^, ATE1 can utilize Arg-conjugated tRNA^Arg^-derived fragments (tRF^Arg^), which are translation-incompetent and thus can potentially serve to shift the balance between translation and arginylation [12]. However, generation of Arg-tRF^Arg^ also requires RARS as the initial step, and thus RARS availability and activity can potentially be rate-limiting for arginylation.

In mammalian cells, RARS exists as two isoforms derived from one mature mRNA [13, 14] via translation from two alternative start codons. Thus, these two RARS isoforms are identical downstream of the second start codon, but the “long” RARS includes an additional N-terminal stretch of sequence containing a leucine zipper (LZ) that scaffolds this RARS into the multi-tRNA synthetase complex (MSC, which contains IARS, LARS, MARS, QARS, RARS, KARS, DARS, EPRS and three scaffold proteins AIMP-1, -2, -3) [15]) largely dedicated to translation [16]. In contrast, “short” RARS, lacking this domain, is soluble and cytosolic. The MSC has been proposed to channel aa-tRNAs directly to ribosomes to support efficient translation [16]. LZ of RARS interacts with AIMP-1, which is required for assembly of RARS in the MSC. This interaction also forms a platform to anchor QARS to the MSC [17]. Although AARSs are exclusively located in the cytoplasm of eukaryotic cells to provide protein synthesis, several studies revealed that MSC-bound AARSs have been also found in the nucleus [17, 18]. It has been previously proposed based on this scaffolding that long RARS functions primarily in translation, while the short RARS could potentially be partially or exclusively dedicated to translation-independent functions, including arginylation [14], however this hypothesis has never been directly tested. It was recently found that displacement of the long RARS from the MSC by deletion of the leucine zipper domain responsible for this scaffolding does not affect global translation levels and tRNA^Arg^ aminoacylation [17], suggesting that the balance between translation and potential translation-independent RARS functions might be more complex.

Here we used in vitro and in vivo assays to investigate the interplay between arginylation, translation, and the availability and composition of RARS isoforms. Our results demonstrate that intracellular arginylation activity does not directly depend on active translation or the levels of RARS enzymes. However, we find that displacement of the long RARS from the MSC increases intracellular arginylation. In the absence of long RARS, ATE1 redistributes into the cytosolic fraction in cells, reminiscent of a similar redistribution of RARS [17]. Our results suggest that ATE1 is linked to non-canonical RARS functions in driving the redistribution of MSC into the cytosol and provide the first comprehensive analysis of the interdependence between translation, arginyl-tRNA^Arg^ synthesis, and arginylation.

## Results

### Intracellular arginylation depends on the physiological state of the cell, but does not compete with active translation

We previously found that arginylation of β-actin, a process linked to the migratory activity of cells, significantly increases in 50% confluent cell cultures, where cells are expected to undergo active division and migration [19]. To test whether changes in cell confluency are accompanied by general changes in arginylation activity, we used the previously developed arginylation sensor – a construct that contains an N-terminal arginylation target peptide fused to GFP that enables ratiometric imaging of arginylation levels using Arg- and GFP-specific antibodies ([20] and Fig 1, left). In these assays, arginylation levels were significantly higher in semi-confluent cells compared to the cells grown to a confluent monolayer (Fig. 1). Thus, physiological state of the cells can affect their arginylation activity.

**Figure 1.**
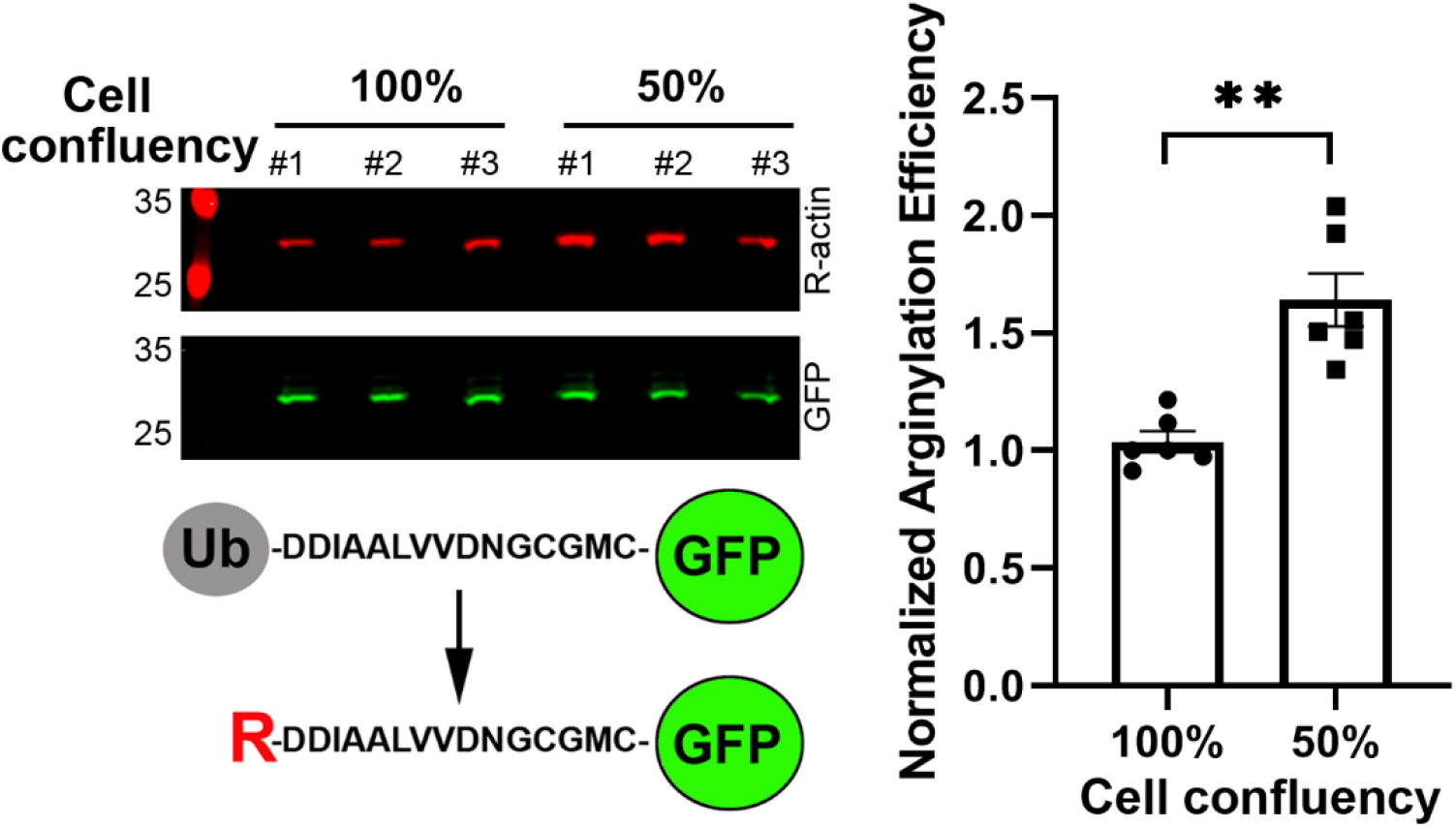
Intracellular arginylation activity depends on the physiological state of the cell. Left, immunoblots (top) and schematic representation (bottom) of the arginylation sensor used in this study to detect intracellular arginylation. In this arginylation sensor, 15-residue peptide based on the β-actin N-terminal sequence is expressed in cells as a fusion with an N-terminal ubiquitin (Ub) moiety and a C-terminal GFP. Arginylation occurs on the N-terminal D of the sequence following cotranslational Ub removal by deubquitinating enzymes and is detected with antibody against the arginylated β-actin (R-actin). Right, quantification of arginylation activity as a ratio of R-actin to GFP signal in human embryonic kidney 293T cells grown to 100% or 50% confluency. Error bars represent SEM, n=6. ** P<0.01, Welch’s *t*-test.

One of the expected changes during cell transition to confluency is a decrease in metabolic activity [21] and translation, a process that utilizes tRNA^Arg^ and RARS activity, which are also required for arginylation by ATE1. To test if arginylation levels change upon translation inhibition, we compared sensor arginylation levels in control cells and cells treated with translation inhibitors cycloheximide and chloramphenicol. We reasoned that if arginylation and translation exist in direct competition, cycloheximide/chloramphenicol treatment should increase arginylation. If, however, arginylation depends on active translation as a donor of reactive compounds, such as Arg-tRNA^Arg^, we should see a decrease in arginylation after cycloheximide/chloramphenicol treatment. However, cycloheximide/chloramphenicol treatment did not change arginylation levels in cultures of similar confluency (Fig. 2A). Thus, inhibition of translation does not appear to affect arginylation.

**Figure 2.**
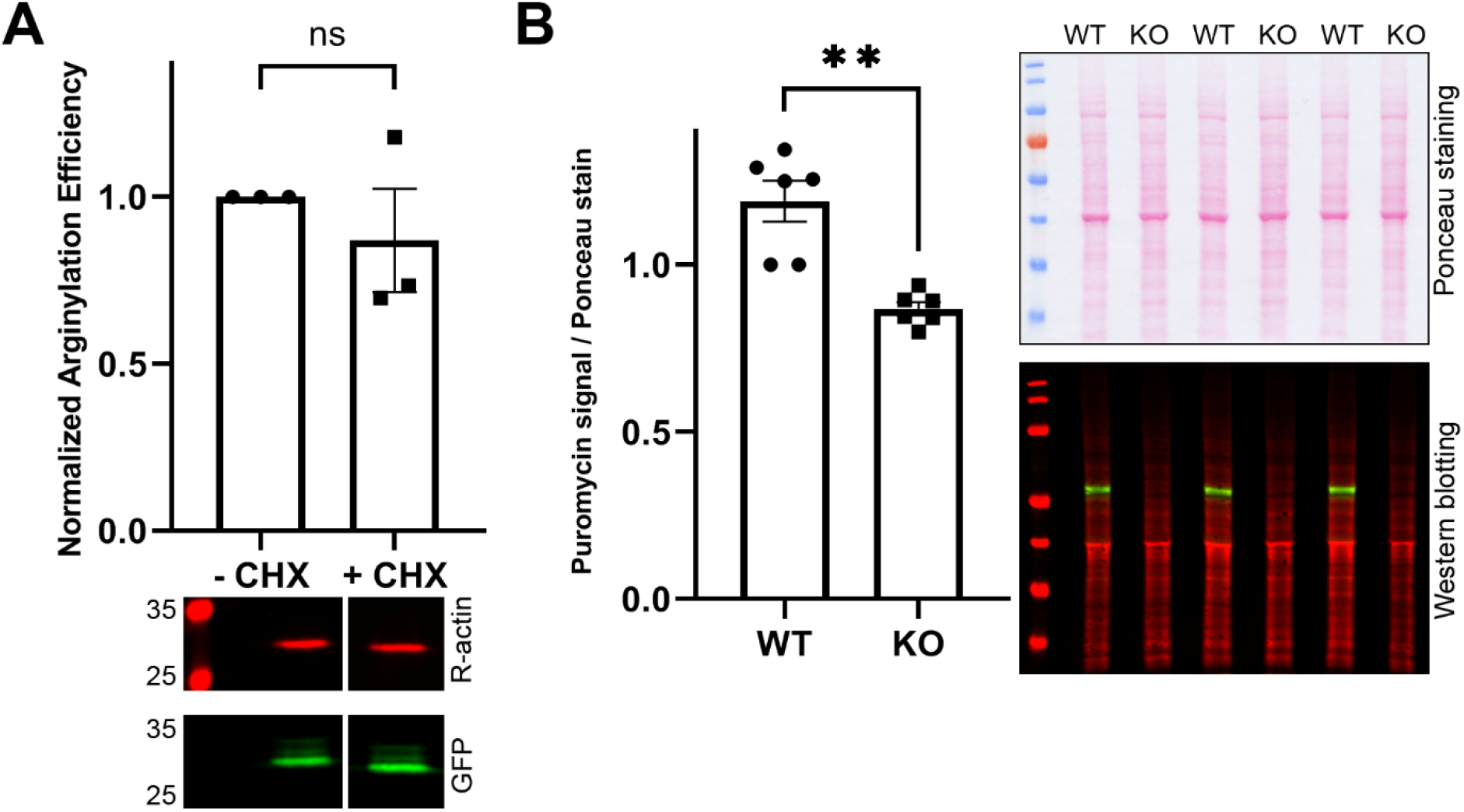
Intracellular arginylation depends on the physiological state of the cell but does not compete with active translation. **A**, Chart (top), and representative immunoblots (bottom) of arginylation sensor quantification in human embryonic kidney 293T cells with and without addition of cycloheximide (CHX) and chloramphenicol. Error bars represent SEM, n=3. ns, not significant, Welch’s *t*-test B, Chart (left), and immunoblots (right) of wild type (WT) and *Ate1* knockout (KO) mouse embryonic fibroblasts stained with Ponceau S for total protein load (top) and puromycin (red) and ATE1 antibodies (green) (bottom). Error bars represent SEM, n=3. ** P<0.01, Welch’s *t*-test.

We also addressed a reciprocal possibility, by testing whether inhibition of arginylation affects translation activity. To do this, we measured overall translation levels in *Ate1* knockout mouse embryonic fibroblasts, previously derived in our lab from *Ate1* knockout mice [9]. As a measure of translation activity, we used puromycin, which mimics the 3’ adenosine of a charged tRNA and incorporates into the C-terminus of elongating nascent chains, thus labeling the entire body of newly synthesized proteins in the cell [22-25]. We reasoned that since *Ate1* knockout cells lack arginylation, if a direct competition between translation and arginylation exists, they should exhibit an increase in translation activity in the absence of ATE1. However, in these assays, overall levels of puromycin staining were significantly lower in *Ate1* knockout cells compared to wild type (Fig, 2B), suggesting that lack of ATE1 in these cells impairs protein synthesis.

Thus, arginylation activity shows an overall dependence on the cells’ physiological state and facilitates translation, but there is no direct competition between translation and arginylation.

### ATE1 can interact with long and short RARS, but overexpression of RARS does not facilitate arginylation

It has been previously proposed in early biochemical studies that ATE1 in vivo is complexed with RARS [26], and that it may functionally interact with short RARS in the cytosol [14], even though direct interaction between ATE1 and RARS has never been demonstrated in any subsequent studies. To test whether ATE1 can directly interact with either of the RARS isoforms, we performed immunoprecipitation using RARS antibodies from cells overexpressing either long or short RARS, generated by editing out either the first or the second translation initiation site in the RARS transcript (Fig. 3A). We then tested these immunoprecipitation steps with antibodies to ATE1. In these assays, ATE1 was present mostly in the input and the flow-through (Fig. 3B, top), however a very minor amount of ATE1 could be detected in both long RARS and short RARS immunoprecipitates (Fig. 3B, bottom). This amount was not stoichiometric to RARS and overall pushed the detection levels by Western blot, suggesting that only a minor amount of ATE1 can potentially be involved in RARS interaction. Thus, the majority of both short and long RARS likely exists in ATE1-free pool. Moreover, the minor ATE1 fraction found in the RARS precipitates does not exhibit any apparent bias toward the long or short RARS isoform.

**Figure 3.**
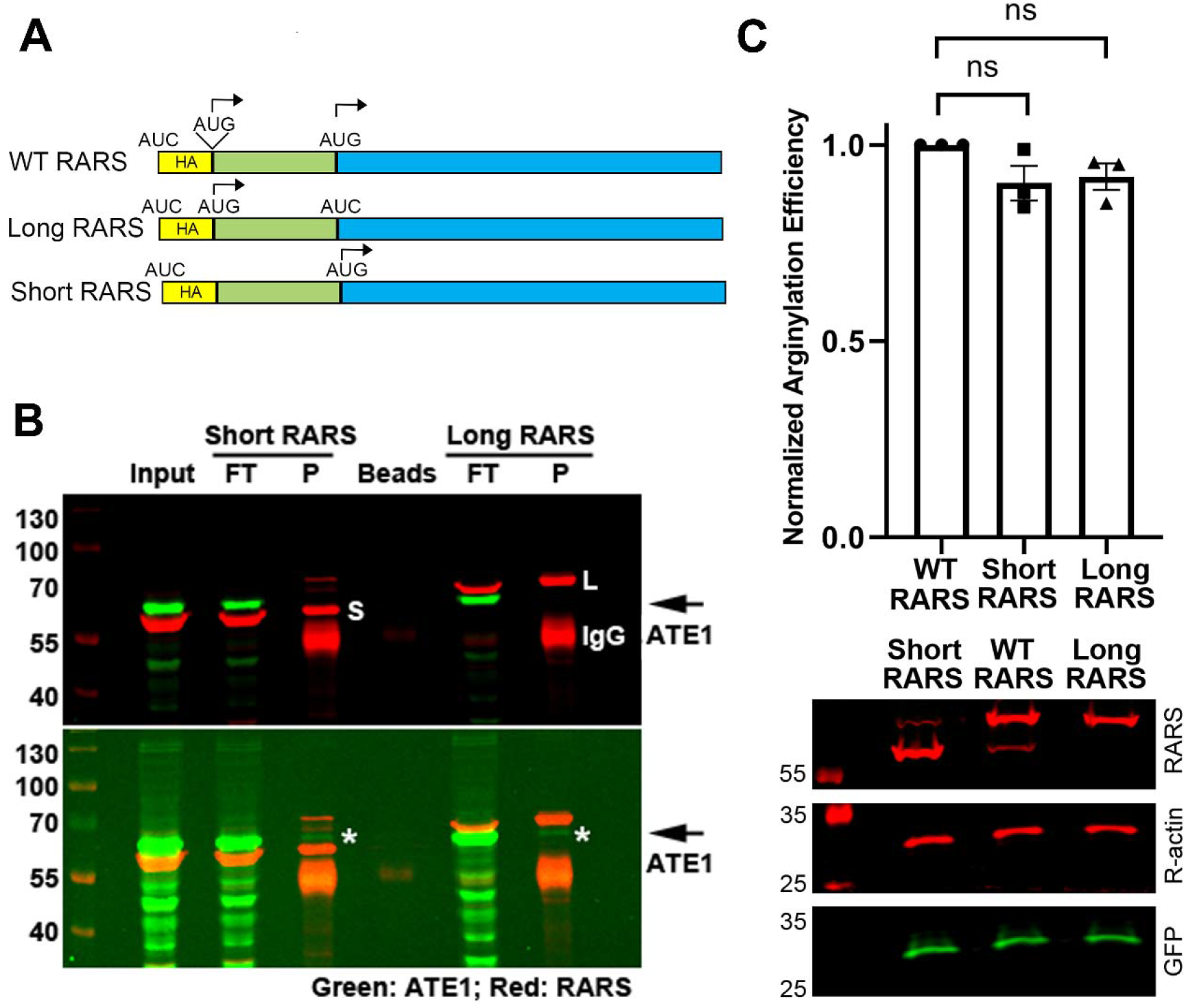
ATE1 can interact with long and short RARS, but overexpression of RARS does not facilitate arginylation. A, Schematic representation of the RARS constructs used for immunoprecipitation and overexpression. Start codons (AUG) were individually mutated to AUC to abolish the HA tag from the expression construct and ensure the start of translation either on both sites (WT), or on the upstream or downstream AUG individually (for the long and short RARS, respectively). B, representative immunoblots of the immunoprecipitation steps from HEK293T cells transfected with either short or long RARS as indicated on the top and visualized with RARS and ATE1 antibodies. FT, flow-through, P, pulldown. Top and bottom panel show the same blot at different exposure to visualize typical immunoprecipitation steps (top), as well as the minor levels of ATE1 in the precipitates (bottom, marked with asterisks and arrows on the right). The position of short and long RARS are indicated in the top panel as S and L, respectively. IgG, immunoglobulin heavy chain. The pulldowns were repeated at least 3 independent times with similar results. C, Chart, and representative immunoblots of arginylation sensor quantification in HEK293T cells transfected with different RARS isoforms as indicated. Error bars represent SEM, n=3. ns, not significant, Welch’s *t*-test.

To test whether increased availability of either short or long RARS can facilitate arginylation, we tested sensor arginylation levels in cells overexpressing the long RARS, the short RARS, or the wild type RARS transcript containing both of the alternative start codons that give rise to both RARS isoforms. There was no difference between arginylation levels in these cells (Fig. 3C). Thus, overexpression of the different isoform of RARS does not facilitate arginylation.

### Displacement of long RARS from the MSC increases intracellular arginylation

To test whether redistribution of RARS between the soluble fraction and the fraction incorporated into the MSC affects arginylation, we used the previously described cell line, in which the leucine zipper in the long RARS has been deleted, thus converting the entire RARS pool into the soluble cytosolic (short) form [17]. In these cells, termed dLZ for delta-leucine-zipper, translation is, surprisingly, not affected, but the entire MSC is restricted from localizing into the nucleus, suggesting that RARS localization via leucine zipper is required for the regulation of the alternative nuclear functions of this enzyme [17].

Comparison of arginylation levels in dLZ and parental wild type cells of similar confluency revealed a significant increase of arginylation in dLZ (Fig. 4A). This result suggests that conversion of intracellular RARS from MSC-bound to cytosolic pool directly or indirectly facilitates arginylation. At the same time, transfection of dLZ cells with either short or long RARS did not affect arginylation (Fig. 4B), consistent with our observation in control cells (Fig. 3C). Together, these results suggest that while displacement of RARS from MCS into the cytosolic pool facilitates arginylation, this effect is not related to RARS enzymatic activity in tRNA charging.

**Figure 4.**
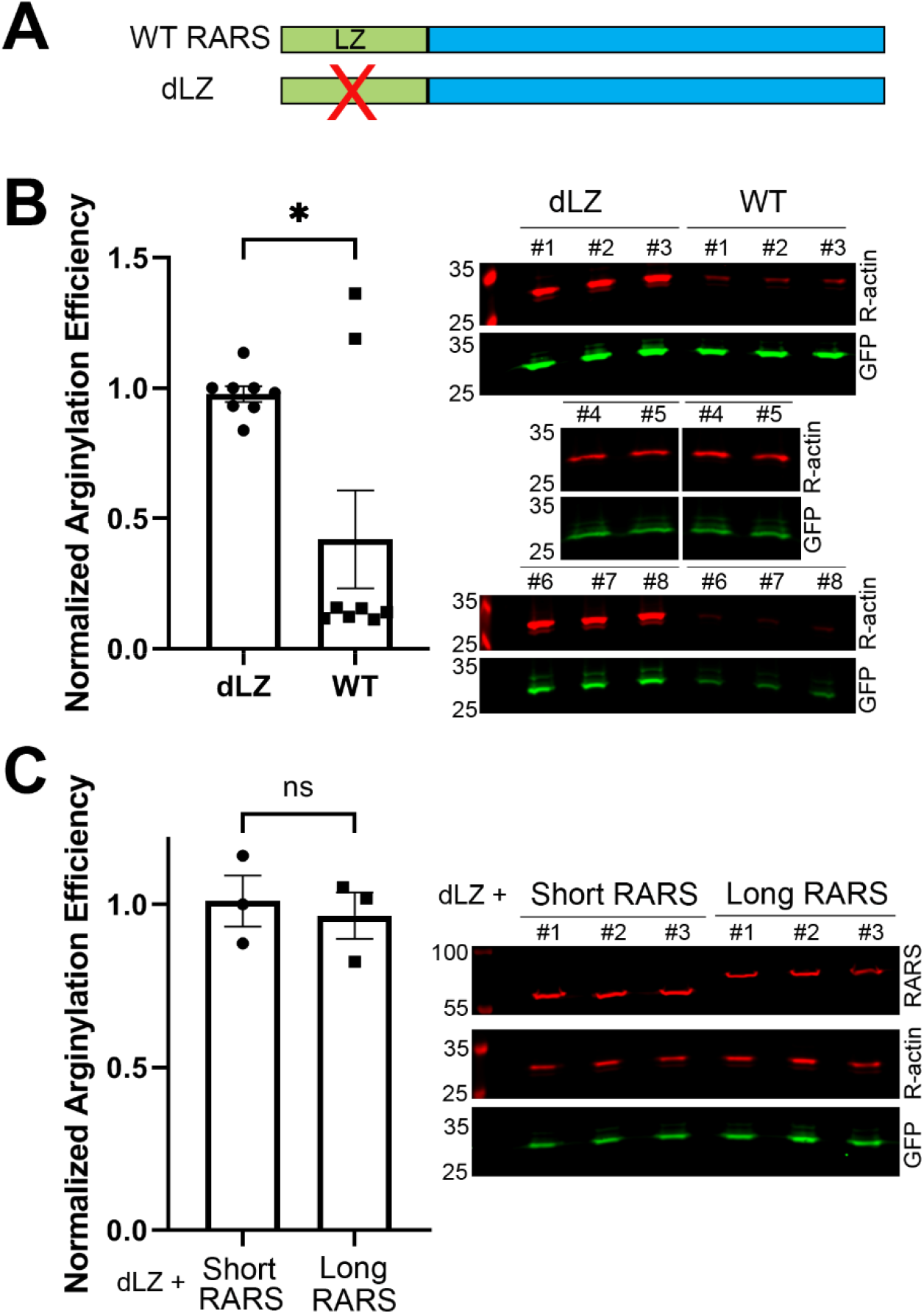
Displacement of long RARS from the multi-synthetase complex increases intracellular arginylation. A, Diagram representing deletion of the leucine zipper (LZ) in dLZ cells to convert the entire RARS into the cytosolic soluble pool. B, Chart, and representative immunoblots of arginylation sensor quantification in HEK293T cells produced by knocking out the leucine zipper domain in long RARS to displace it from the multi-synthetase complex (dLZ) compared to the parental cell line (WT). Cells of similar confluency were chosen as pairs for quantification. Error bars represent SEM, n=8. * P<0.05, Welch’s *t*-test. C, Chart, and representative immunoblots of arginylation sensor quantification in dLZ cells transfected with long or short RARS. Error bars represent SEM, n=3. ns, not significant, Welch’s *t*-test.

### Displacement of RARS from the MSC increases the cytosolic fraction of ATE1

In dLZ cells, the only effect on RARS found in the previous study was independent of its enzymatic activity and its role in translation, but linked to its non-canonical role in facilitating localization of MSC to the nucleus [17]. Two of the four ATE1 isoforms has been previously shown to exhibit partial, transient nuclear localization [11, 27, 28]. Making a parallel with the RARS, it is conceivable that deletion of the leucine zipper and the ensuing displacement of AARSs from the nucleus may also affect the nuclear:cytosolic balance of ATE1. To test this, we compared the levels of ATE1 in the cytosolic fraction in dLZ cells and control. Remarkably, ATE1 cytosolic fraction was significantly increased in dLZ cells (Fig. 5A), while overall ATE1 level in these cells was not changed (Fig. 5B). Thus, displacement of RARS from the MSC facilitates ATE1 redistribution into the cytosol. We speculate that this change likely leads to the increased arginylation in dLZ cells (Fig. 4A), since the arginylation sensor we are using is expected to be cytosolic.

**Figure 5.**
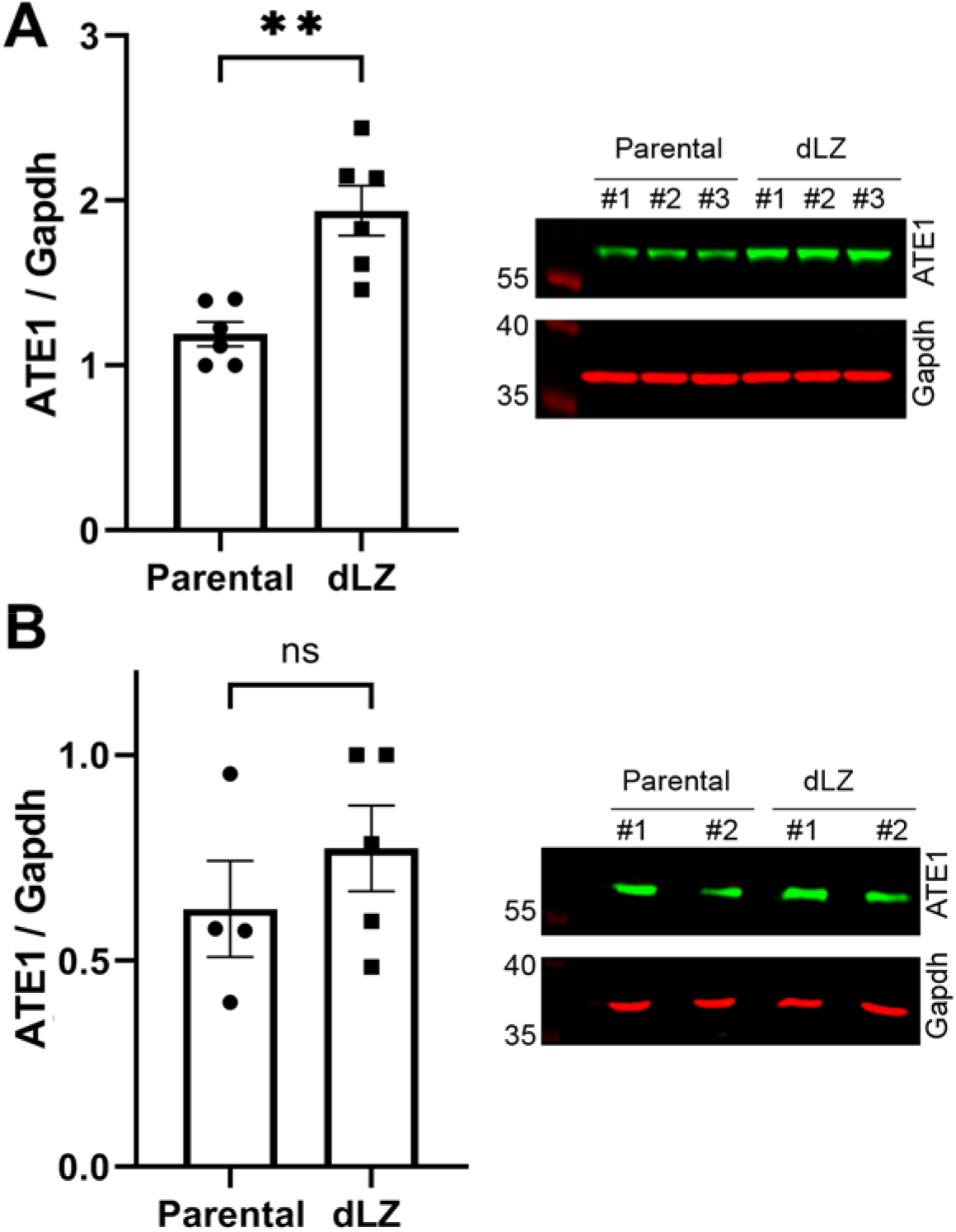
Displacement of long RARS from the multi-synthetase complex induces ATE1 redistribution into the cytosol. A, Chart and representative immunoblots of ATE1 levels in the cytosol of wild type (parental) and dLZ cells. B, Chart and representative immunoblots of total ATE1 levels in wild type and dLZ cells. Error bars represent SEM, n=3. ** P<0.01, Welch’s t-test.

## Discussion

This work represents the first study of the functional interplay between arginylation, translation activity, and RARS enzymes. We find that arginylation does not directly compete with the translation machinery and is not directly dependent on intracellular RARS levels, but is functionally linked to RARS in a translation-independent manner, potentially facilitated by direct or indirect ATE1-RARS interaction (Fig. 6). Identification of the components of this interaction constitutes an exciting direction of future studies.

**Figure 6.**
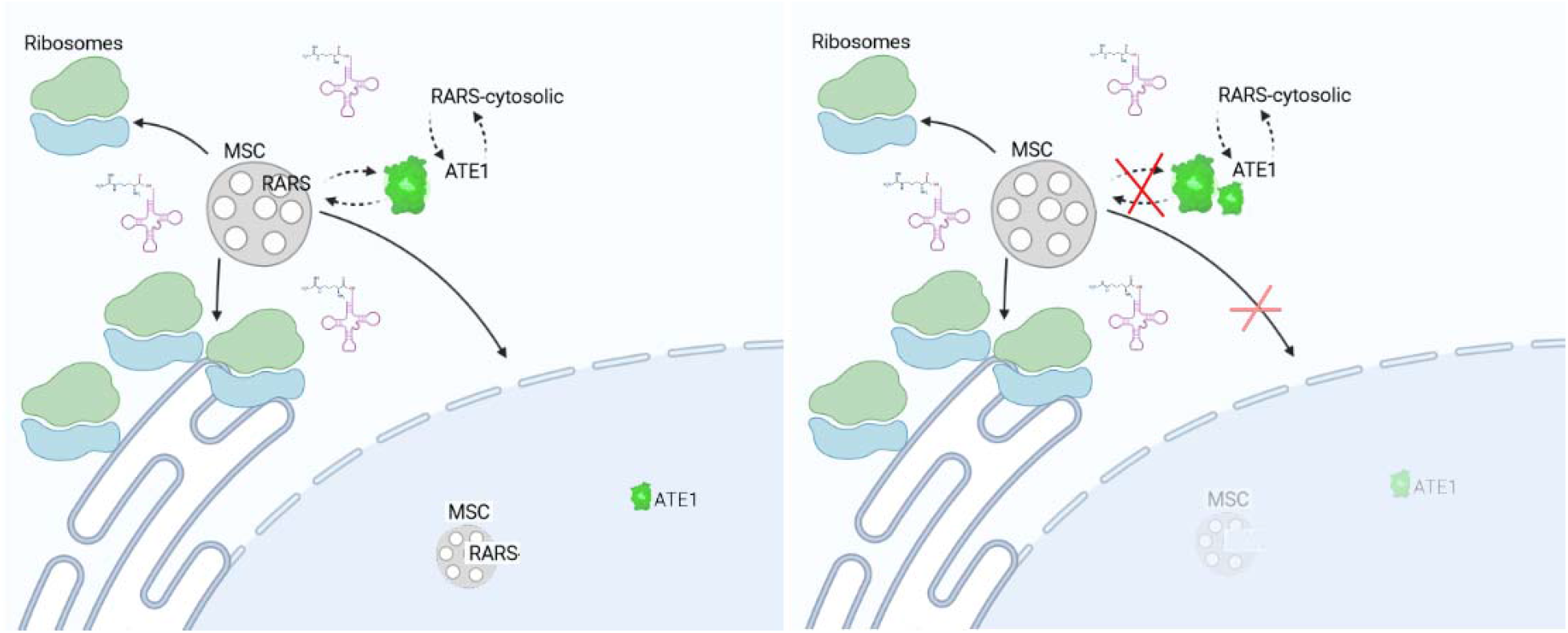
Interplay between ATE1 and RARS. In wild type cells (left) ATE1 exists in a balance with cytosolic and MSC-bound RARS, as well as their nuclear fractions. Knockout of the leucine zipper (right) that scaffolds long RARS to the MSC perturbs this balance by reducing the nuclear MSC and ATE1 localization, resulting in an increase of cytoplasmic ATE1 activity. Cytosolic and nuclear environments are denoted with white and blue background, respectively.

Our data reveal that intracellular arginylation activity depends on physiological state of the cell, and is higher in semi-confluent cells active in the cell cycle, compared to the resting cells in the dense monolayer. This finding is in agreement with our previously published data demonstrating a similar increase in β-actin arginylation in 50% confluent cell cultures, compared to cells grown to 100% confluency [19]. While the specific factors driving this difference remain to be identified, this finding strongly suggests the existence of upstream mechanisms that regulate arginylation at the global cellular level. Investigating the contribution of individual pathways to this regulation will shed light on the overall role of arginylation.

ATE1 has been previously proposed to act in direct competition to the translation machinery that utilized the same molecule, Arg-conjugated tRNA^Arg^, as the ribosome [12]. Our data show that inhibition of translation does not affect intracellular arginylation levels. Notably, cycloheximide/chloramphenicol treatment has been used in multiple prior studies to suppress incorporation of Arg into proteins during translation and identify specifically proteins that incorporate Arg through posttranslational arginylation (see, e.g., [7, 29]), however these studies never addressed whether cycloheximide treatment changes arginylation levels in cells. Our current result strongly suggests that there is no direct competition between translation and arginylation. Since most of tRNA^Arg^ in cells are charged with Arg to over 90% [30], it is possible that they are sufficiently abundant for both pathways. It is also possible that ATE1 depends largely on translation-incompetent Arg-tRF^Arg^, previously demonstrated to be efficient donors of the Arg group for arginylation [12].

We find that lack of ATE1 leads to overall lower translation activity in cells. Given our data suggesting that ATE1 does not directly compete with protein synthesis, it seems likely that this effect is due to other mechanisms that may regulate translation through arginylation. It is possible, for instance, that the components of the translation machinery are directly arginylated, and that lack of this regulation in *Ate1* knockout cells can directly inhibit the activity of these components. This possibility constitutes an exciting direction of future studies.

To date, very few ATE1 interacting proteins have been identified. Our finding that a fraction of ATE1 can bind to RARS adds an important interaction partner to this very short list. In principle, RARS binding can ensure that ATE1 is strategically placed to utilize newly conjugated arginyl-tRNA^Arg^ before they can be used for translation. Even though our study suggests that RARS availability is not rate-limiting for arginylation, this mechanism can potentially contribute to regulation of arginylation in vivo.

Given the extremely minor levels of ATE1 found in RARS pulldowns, it is possible that additional unknown players or interactors can participate in the interplay of arginylation and translation. A hypothesis-free discovery approach, such as interaction proteomics, may provide novel candidates for future studies.

Our data suggest that disruption of the MSC can affect ATE1’s redistribution into the cytosol. Previously two of the four ATE1 isoforms in mammalian cells have been found to exhibit transient localization in the nucleus [11, 27, 28]. A small fraction of ATE1 has also been found to localize to the mitochondria [31]. Since displacement of RARS from MSC in dLZ cells does not affect total ATE1 levels, it is likely that ATE1’s cytosolic increase in these cells occurs due to its redistribution from either the nuclear and/or the mitochondrial fraction into the cytosol. Given that RARS scaffolds components of the MSC into the nucleus, it is attractive to suggest that it also participates in ATE1’s nuclear shuttling, so that disruption of the RARS nuclear localization would also affect ATE1. This idea is in line with the fact that a fraction of ATE1 can co-immunoprecipitate with RARS. Going further in this reasoning, it is possible that ATE1 may be directly or indirectly linked to the alternative function of RARS in RNA editing, and potentially other processes in the nucleus. This possibility, and the potential interlace of ATE1 and RARS’s non-canonical functions constitute an exciting direction of future studies.

## Materials and Methods

### Materials

Human Embryonic Kidney 293T (HEK293T) parental cells and cells with the deletion of the Leu zipper domain in the long RARS (dLZ) [17] were a generous gift from Dr. Paul Schimmel (the Scripps Research Institute). Immortalized wild type and *Ate1* knockout mouse embryonic fibroblasts (MEF) cells obtained in the lab as described in [8]. Cells were grown in Dulbecco’s modified Eagle’s medium with GlutaMAX™ supplement (DMEM + GlutaMAX, Gibco) with 10% fetal bovine serum and 1% Penicillin-Streptomycin at 37°C with 5% CO_2_.

Arginylation sensor plasmids were a generous gift from Dr. Fangliang Zhang (University of Miami). Plasmids expressing long and short RARS were generated from a commercial RARS plasmid (HG19710-NY, Sino Biological) by site-directed mutagenesis in our laboratory.

The following antibodies were used in this study: rabbit anti-R-actin (ABT264 EMD Millipore, 1:2000), mouse anti-GFP (Ab1218 Abcam, 1:3000), rabbit anti-RARS (orb247357 Biorbyt, https://www.biorbyt.com/rars-antibody-orb247357.html, 1:5000), mouse anti-puromycin (MABE343 Millipore Sigma, 1:3000), rat anti-ATE1 (homemade, 1:1000) or mouse anti-GAPDH (Ab8245 Abcam, 1:5000).

All the chemicals used for cell treatment, including cycloheximide, puromycin, and chloramphenicol, were obtained from Sigma-Aldrich.

### Total Protein Analysis and Cell Fractionation

For total protein analysis, cells were washed once with PBS and harvested by scraping and centrifugation. Cell pellets were lysed in lysis buffer (50 mM Tris-HCl, pH 7.5, 150 mM NaCl, 0.1% Triton X-100, with protease inhibitor cocktail (Sigma-Aldrich)) at the w:v ratio of cell pellet:lysis buffer 1:5, followed by vortexing for 5-10 min and sonication (GE Healthcare) on ice at level 5 for 20 times (3 s pulse 3 s pause). Cell lysates were clarified by centrifugation at 16,000 x g for 15 min at 4°C. Protein concentration was measured by Pierce™ BCA Protein Assay Kit.

Cell fractionation into nuclear and cytosolic fraction was performed as described in [17] with some modifications. 5 × 10^6^ HEK293T cells were seeded on a 10 cm dish one day before the procedure. The cells were washed once with PBS and harvested by scraping and centrifugation. The cell pellet was incubated at -80°C for 45 min and dissolved into 250 μl cell fractionation buffer (20 mM HEPES, pH 7.5, 10 mM KCl, 1 mM MgCl_2_, 1 mM EDTA, 1 mM DDT, complete protease inhibitors). The cells were lysed by passing through a 27 g needle for 20 times and vortexing for 30 s. The sample was centrifuged at 800 x g for 5 min at 4°C. The supernatant was taken carefully and centrifuged at further 16,000 x g for 15 min at 4°C. The supernatant was mixed with 4 x SDS loading buffer, followed by boiling for 10 min. 10 μl of cytoplasmic fraction was loaded for SDS-PAGE electrophoresis.

### Cell treatments for arginylation sensor measurements, translation inhibition and puromycylation assay

Cell transfections were performed in 6-well plates using Lipofectamine 2000 (Thermo Fisher Scientific) according to the manufacturer’s protocol, using 1:3 ratio of DNA:Lipofectamine (4 μg DNA: 12 μl Lipofectamine).

For arginylation sensor experiments, cells were transfected with arginylation sensor plasmid 24-48 hours prior to the experiment. For arginylation measurements in confluent and semiconfluent cultures shown in Fig.1, cells transfected with the sensor were split after 24 h to achieve 50% and 100% confluency and incubated for further 24 h in culture. For Fig.2A, 48 h after transfection, 100 μg/mL cycloheximide and 40 μg/mL chloramphenicol were added to the culture media, followed by further incubation for 1 h. For Fig.3C, equimolar mixture of arginylation sensor and RARS plasmids were transfected into the cells 48 h prior to harvesting.

On the day of the experiment, the transfected cells were washed once with phosphate-buffered saline (PBS, Corning) and harvested by scraping and centrifugation. Cell pellets were lysed in 4 x SDS loading buffer at the w:v ratio of 1:20 (1 mg cell pellet: 20 μl buffer), followed by boiling the samples for 10 min. 10 μl of each sample was loaded for SDS-PAGE electrophoresis and analyzed by Western blot.

For puromycylation assay, wild type and *Ate1*^-/-^ MEFs were treated with 10 μg/mL puromycin for 15 min at 37°C. Cells were collected and lysed by sonication to measure total protein concentration using Pierce™ BCA Protein Assay Kit. 16 μg total protein/lane of each sample was loaded for SDS-PAGE electrophoresis.

### Immunoprecipitation

For anti-RARS immunoprecipitations (IP), dLZ or parental HEK293T cells were grown in 10 cm dishes to 60-80% confluency and transfected with plasmids expressing short, long, or wild type RARS as indicated in the figures. 48 hours after the transfection, cells were harvested into lysis buffer and sonicated as described above in the “total protein analysis” section. 50 μl of Protein A agarose beads (Invitrogen) pre-equilibrated with the lysis buffer were added to clear up the lysate for 1 h at room temperature on a rocker. Following removal of the beads by centrifugation, 10 μg of anti-RARS antibody (orb247357 Biorbyt) was added to the cell lysate and incubated on a rocker for 1 h at room temperature, followed by addition of 50 μl of Protein A agarose beads (Invitrogen) pre-equilibrated in the lysis buffer and additional incubation for 1 h at room temperature on a rocker. The beads were collected by centrifugation and washed for three times with the lysis buffer. 20 μl of 4 x SDS loading buffer was added to the beads and boiled, followed by analysis by SDS-PAGE electrophoresis and Western blotting.

### Western blotting

The gels were transferred to nitrocellulose membrane at 100 V for 60 min. The blots were then blocked by 5% milk in PBS-Tween at 4°C for 16 h and incubated with primary antibodies for 60 min at room temperature, followed by washes and incubation with secondary antibodies (1:5000) conjugated to IRDye800 or IRDye680. Images were acquired and analyzed by Odyssey Imaging System (LI-COR).

## Acknowledgements

We thank members of the Kashina lab for helpful discussions. This work was supported by the NIH grant R35GM122505 to AK.

## Author contributions

IA and AK, designed experiments, analyzed data, wrote paper

IA, performed experiments

AK, supervised the project

